# Structuring Role of Tau-Tubulin Co-Condensates in Early Microtubule Organization

**DOI:** 10.1101/2024.11.04.621946

**Authors:** Chaelin Lee-Eom, Jaehun Jung, Celine Park, Min Ju Shon

## Abstract

Tau protein, a key microtubule-associated protein in neurons, is traditionally known for stabilizing microtubules. However, its recently discovered ability to undergo liquid-liquid phase separation (LLPS) reveals a broader, dynamic role in nucleating and organizing microtubule networks. Here, using a combination of real-time imaging and a geometric approach based on Voronoi tessellation, we examined how tau condensation leads to clustering and local tubulin enrichment, supporting microtubule organization. Our observations show that tau-tubulin co-condensates not only initiate nucleation and branching of microtubules but also drive the network’s gradual evolution through a “dynamic weaving” process. By generating Voronoi diagrams from super-resolution and confocal microscopy images of the stabilized network, we quantitatively mapped tubulin enrichment as a function of tau density, revealing that high-density tau clusters, approximately 0.2 μm in size, correlate with tubulin-rich spots at equilibrium. Overall, these findings provide new insights into tau-tubulin co-condensates as dynamic structuring elements, whose liquid-like properties continuously reshape the microtubule network, creating a flexible and adaptive architecture essential for cellular function.

## Introduction

Microtubules are dynamic cytoskeletal filaments essential for cellular functions like maintaining cell shape, intracellular transport, and mitosis. Built from tubulin dimers, microtubules exhibit dynamic instability, alternating between polymerization and depolymerization phases in response to cellular signals, allowing them to perform diverse roles across different cell types^1–3^. For instance, in neurons, microtubules facilitate neurite extension and axonal transport, highlighting their adaptability to specialized cellular environments^4–6^.

The stability and organization of microtubules are tightly regulated by microtubule-associated proteins (MAPs), among which tau plays a particularly crucial role^7^. Traditionally, tau is regarded as a stabilizer, binding along tubulin monomers to reduce depolymerization rates and promoting bundling to resist mechanical stress and enzymatic severing^8–10^. However, recent studies reveal a more dynamic role for tau, especially through its capacity for liquid-liquid phase separation (LLPS), forming condensates that interact with tubulin and nucleic acids. These tau condensates recruit and concentrate tubulin within their clusters, facilitating localized nucleation and elongation of microtubule networks^11^. Such properties suggest tau’s role extends beyond stabilization, positioning it as an architect of the microtubule network through mechanisms like nucleation, branching, and structural remodeling.

Understanding tau’s involvement in microtubule dynamics is especially relevant to neuronal development and health, as aberrant tau-tubulin interactions are implicated in neurodegenerative diseases such as Alzheimer’s^12–15^. In vitro studies, often conducted with supra-physiological tau and tubulin concentrations and molecular crowding agents, have yielded valuable insights into tau’s stabilizing function^11,16–19^. Yet, translating these findings to cellular contexts remains challenging, especially in terms of tau’s structural contributions during early microtubule assembly in crowded intracellular environments. Despite extensive research into tau’s biochemical role in microtubule stabilization, the structural organization of tau-microtubule networks, particularly in early assembly phases, is not yet fully understood. A systematic approach is needed to quantify these networks’ structural properties and clarify tau condensates’ organizational role, facilitating better translation of in vitro findings to in vivo systems.

In this study, we investigate the structuring role of tau-tubulin co-condensates in early microtubule organization, using advanced imaging techniques such as direct stochastic optical reconstruction microscopy (dSTORM), 3D confocal microscopy, and total internal reflection fluorescence (TIRF) microscopy. To analyze the spatial relationships between tau condensates and tubulin, we applied Voronoi tessellation, a mathematical framework that partitions space into regions based on proximity to specific points, recently applied to biophysical structures^20–23^. This approach reveals how tau clusters guide tubulin enrichment dynamics and microtubule formation. Our results show that tau-tubulin co-condensates not only initiate microtubule nucleation but also actively participate in assembling and dynamically organizing the growing network through liquid-like behaviors, including gliding, fusion, and remodeling. This study expands the understanding of tau’s role in cytoskeletal organization, proposing a model in which tau-tubulin co-condensates function as dynamic structuring elements, continuously weaving and adapting the microtubule network.

## Results

### In vitro microtubule polymerization with physiological concentrations of tau

Previous in vitro studies using high concentrations of tau protein (above 10 μM) have demonstrated that tau can nucleate and stabilize microtubules. We aimed to investigate whether similar effects occur at tau concentrations closer to physiological levels (Fig. 1). To this end, we added varying concentrations of tau (0 to 25 μM) to a solution containing 5 μM tubulin and 1 mM GTP to enable microtubule polymerization. For fluorescence imaging, a fraction of the tau and tubulin molecules were labeled with Alexa 488 and HiLyte 647, respectively. The tubulin concentration approximated its intracellular level as reported in previous studies^16,24^, and 5% PEG was included to mimic the macromolecular crowding conditions within cells.

**Figure 1.**
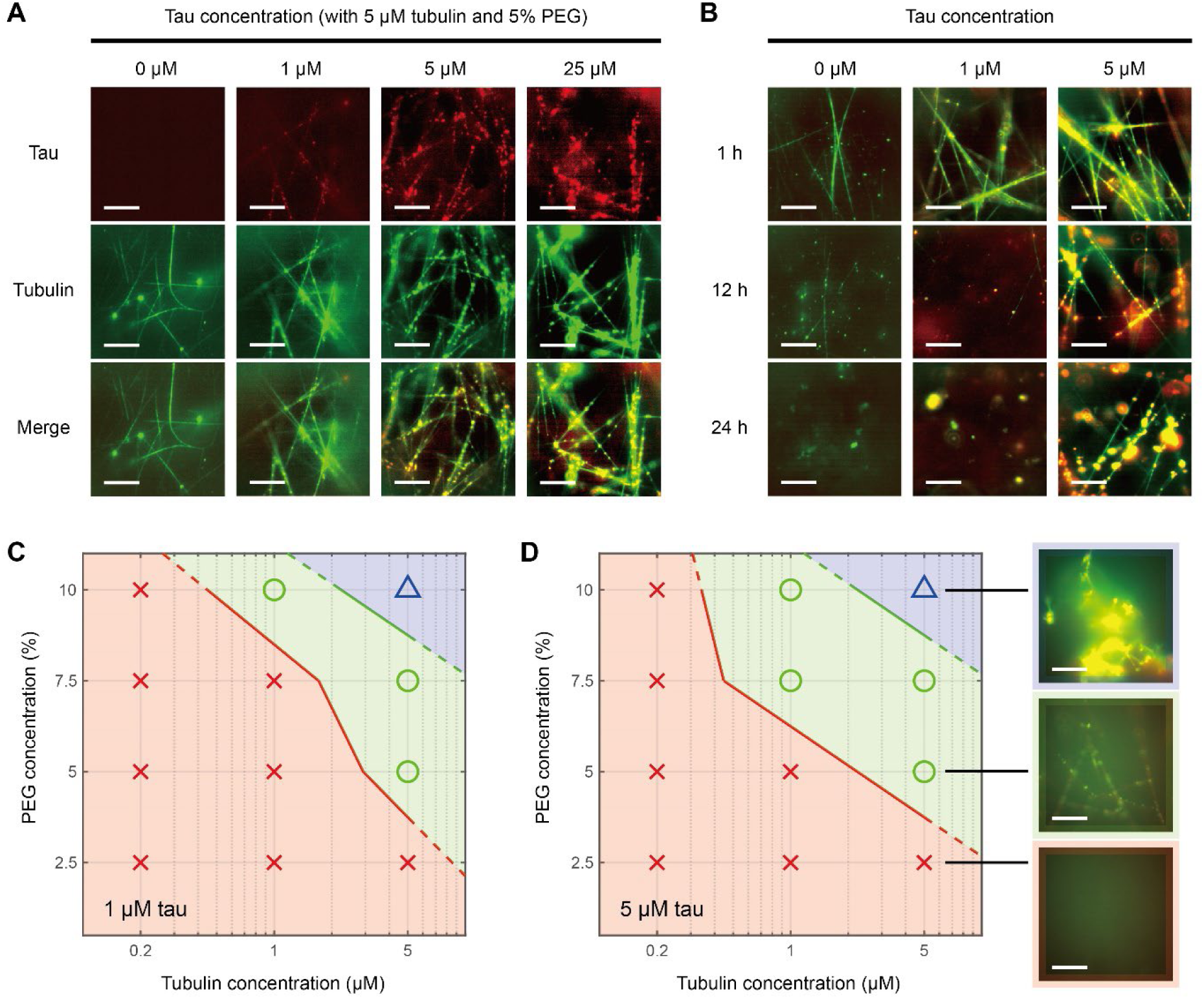
Systematic evaluation of microtubule polymerization in a tau-tubulin-PEG system. (A, B) Fluorescent images of tau and tubulin at varying tau concentrations, with 5% PEG and 5 μM tubulin held constant. In (B), microtubule stability was monitored over 24 h. (C, D) Phase diagrams showing the effects of tau, tubulin, and PEG concentrations on microtubule polymerization. Red crosses indicate conditions where polymerization failed, green circles show successful microtubule formation, and the blue triangle represents conditions with amorphous clumps or aggregates instead of stable polymerization. Insets display representative images of tau and tubulin structures under each condition.

Microtubule filaments assembled immediately after preparing the solutions under all tested conditions (Fig. 1A). While a small number of microtubules formed even without tau, increasing tau concentrations led to a denser and more intricate microtubule network. The structures exhibited a strong correlation between tau and tubulin fluorescence, indicating a direct role of tau in promoting microtubule growth. Notably, at tau concentrations above 5 μM, we frequently observed fluorescent puncta containing both tau and tubulin, consistent with the known co-condensation behavior of these proteins. These findings suggest that micromolar concentrations of tau are sufficient to promote microtubule assembly in crowded solutions, primarily by forming local tau-tubulin co-condensates.

To evaluate the stability of the microtubules formed under these conditions, we monitored their integrity over 24 hours at various tau concentrations (Fig. 1B). Microtubules assembled with 0– 1 μM tau began to depolymerize within a few hours and had completely disappeared after 24 hours. In contrast, those formed with 5 μM tau remained stable for at least 12 hours. Although microtubules assembled at higher tau concentrations eventually disassembled—consistent with intrinsic dynamics following GTP hydrolysis—the tau and tubulin remained colocalized even after microtubule breakdown, leaving behind small clusters (Fig. 1B, 5 μM tau at 24 h). These observations suggest that the strong interaction between tau and tubulin promotes local tubulin enrichment, allowing it to reach the critical concentration required for microtubule nucleation even at moderate tau levels. Moreover, this interaction stabilizes the assembled microtubules over extended periods.

To systematically investigate microtubule polymerization in the tau/tubulin/PEG mixture, we fixed the tau concentration at either 1 μM or 5 μM and varied the concentrations of tubulin and PEG (Fig. 1C, D). Our findings revealed an optimal window of tubulin and PEG concentrations that yielded well-formed microtubule filaments. Low concentrations of tubulin or PEG were insufficient to drive microtubule assembly, whereas excessively high concentrations resulted in amorphous clumps of tau and tubulin molecules (Fig. 1D, inset). The resulting phase diagrams demonstrate that successful microtubule assembly requires a delicate balance among tau, tubulin, and PEG. Any imbalance between tau and tubulin, excessive crowding leading to aggregation, or kinetic trapping during rapid nucleation can impair polymerization dynamics.

### Tau-tubulin co-condensates in nucleating microtubules

While tau-tubulin co-condensates have been shown to promote microtubule growth, we hypothesized that these clusters might cooperatively organize the spatiotemporal development of the microtubule network. To investigate this, we focused on the role of tau during the early phase of microtubule assembly. In our previous experiments using PEG as a molecular crowding agent, microtubule polymerization occurred extremely rapidly, limiting our ability to resolve early events. Interestingly, substituting dextran for PEG as the crowding agent significantly slowed microtubule assembly, allowing us to monitor network formation over the course of an hour (Fig. 2A, Movie S1). The mechanisms underlying this difference require further study, but may involve distinct effects such as protein condensation, weak hydrophobic interactions, and steric exclusion. Based on this observation, we transitioned to a tau/tubulin/dextran system for kinetic studies of microtubule nucleation dynamics.

**Figure 2.**
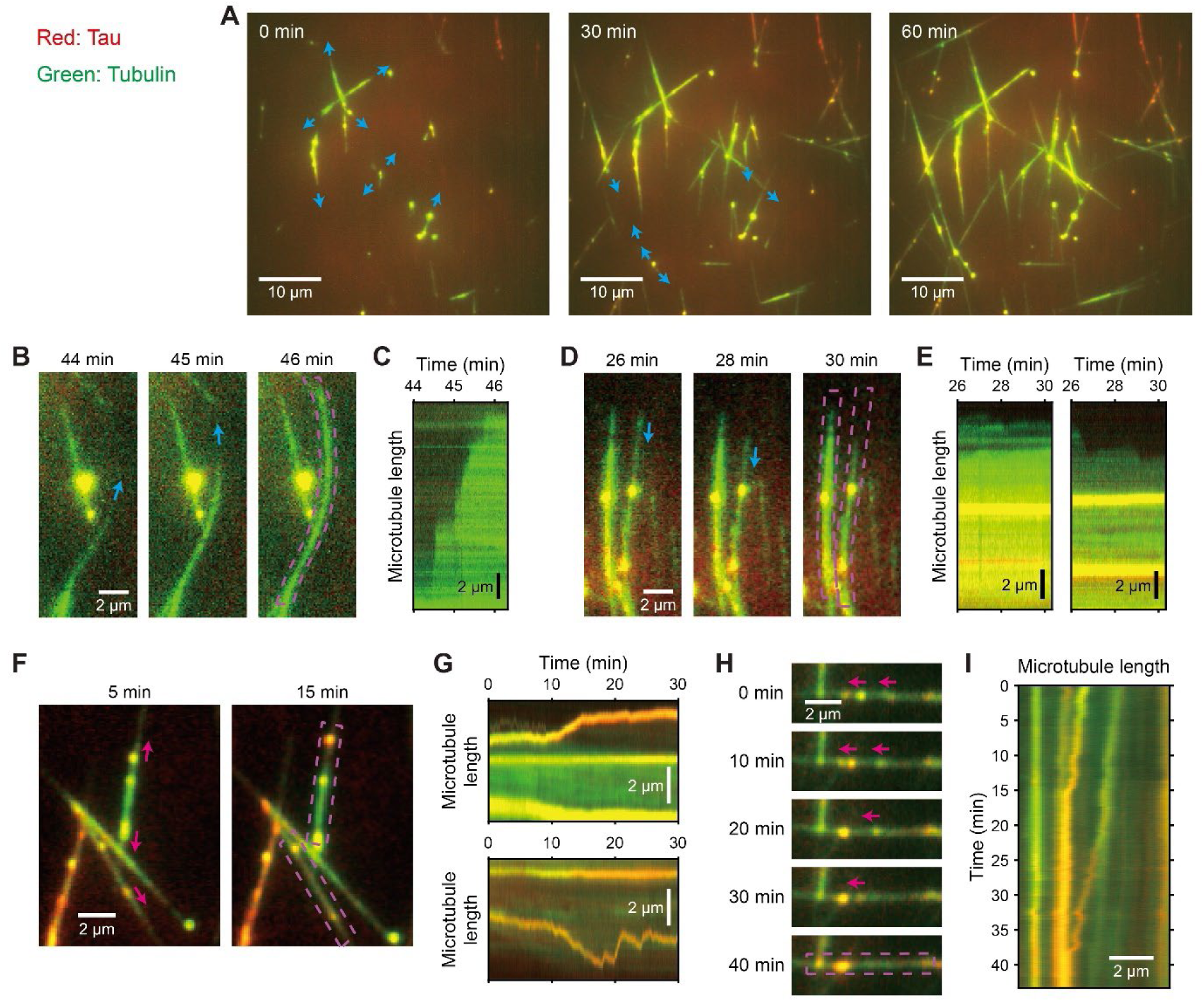
Dynamic nucleation of microtubules by tau-tubulin co-condensates. (A) Time-lapse images showing tau-tubulin co-condensates facilitating microtubule nucleation and extension, with blue arrows indicating sites of microtubule growth over time. (B–E) Sequential images (B, D) and corresponding kymographs (C, E) illustrating changes in microtubule length, with blue arrows marking the direction of extension. Panels (D) and (E) highlight variability in microtubule stabilization by tau-tubulin co-condensates. (F–I) Sequential images (F, H) and kymographs (G, I) showing tau-tubulin co-condensate movement along microtubules, with red arrows indicating the direction of migration. Panels (H) and (I) display merging events between tau-tubulin co-condensates. In all cases, pink dashed boxes mark the regions analyzed for the kymographs in the following panel.

Although overall microtubule assembly continued for over an hour, individual microtubules grew relatively quickly (a few micrometers per minute) (Fig. 2B, C, Movie S2), consistent with timescales reported in previous studies. Intriguingly, we observed that small tau-tubulin co-condensates acted as nucleation sites, promoting bursts of microtubule growth at specific locations (Fig. 2A, blue arrows). Additionally, the brightness of the microtubule assemblies was highly heterogeneous (Fig. 2A–D), suggesting that varying numbers of microtubule filaments were bundled within each segment. Consistent with tau’s stabilizing role, the nucleation of new microtubules was swiftly followed by the spreading of tau onto the surfaces of the newly synthesized filaments, as indicated by the overlap of tau and tubulin fluorescence (Fig. 2A, red and green). In these cases, the fluorescence intensities of tau and tubulin correlated, implying that increasing amounts of tau on microtubules further stabilize them by bundling multiple nascent filaments together. However, not all microtubules were equally stabilized by tau (Fig. 2D, E, Movie S3). We observed that microtubules less supported by the condensates underwent cycles of shortening and regrowth, suggesting that the dynamic instability of microtubules remains a crucial mechanism for shaping the network. These results reinforce the active role of tau-tubulin co-condensates in the spatiotemporal organization of the microtubule network.

### Dynamic nature of tau-tubulin co-condensates

Given the significant influence of tau-tubulin co-condensates on microtubule organization, we questioned whether the resulting reticular geometry is deterministic and static, based solely on the initial locations of these co-condensates. To explore this, we examined the mobility of tau-tubulin co-condensates along the microtubules. Notably, these co-condensates exhibited considerable diffusivity, especially when the filaments were coated with tau and tubulin (Fig. 2F, Movie S4). The corresponding kymographs displayed dynamic disorder during movement, occurring on timescales similar to temporal changes in microtubule dynamics (Fig. 2C, E), suggesting a coupling between condensate diffusion and microtubule behavior.

Furthermore, consistent with their liquid-like nature, the co-condensates often merged into larger clusters by fusing upon collision (Fig. 2H, I, Movie S5). It is possible that the rapid unidirectional movements correspond to microtubule growth and shrinkage, while the slower bidirectional movements represent one-dimensional free diffusion along the microtubules, decoupled from their dynamic remodeling. These observations suggest that tau-tubulin co-condensates do not impose a static, predetermined geometry but rather allow the network to be continuously reshaped through their mobility and interactions.

### Combined effects of co-condensates on the weaving of microtubule network

Building upon the above findings, we explored how the combined actions of tau-tubulin co-condensates may contribute to the dynamic remodeling of the microtubule network. We observed that the condensates actively facilitate both the longitudinal joining and lateral crosslinking of microtubules, promoting linear extension and branching of the network. By bringing microtubule ends together and enabling their alignment and bundling (Fig. 3A, B, Movie S6), they extend structures into continuous filaments. They also mediate lateral crosslinking (Fig. 3C, D, Movie S7), forming stable cross-shaped junctions. These activities enhance the structural integrity and flexibility of the network.

**Figure 3.**
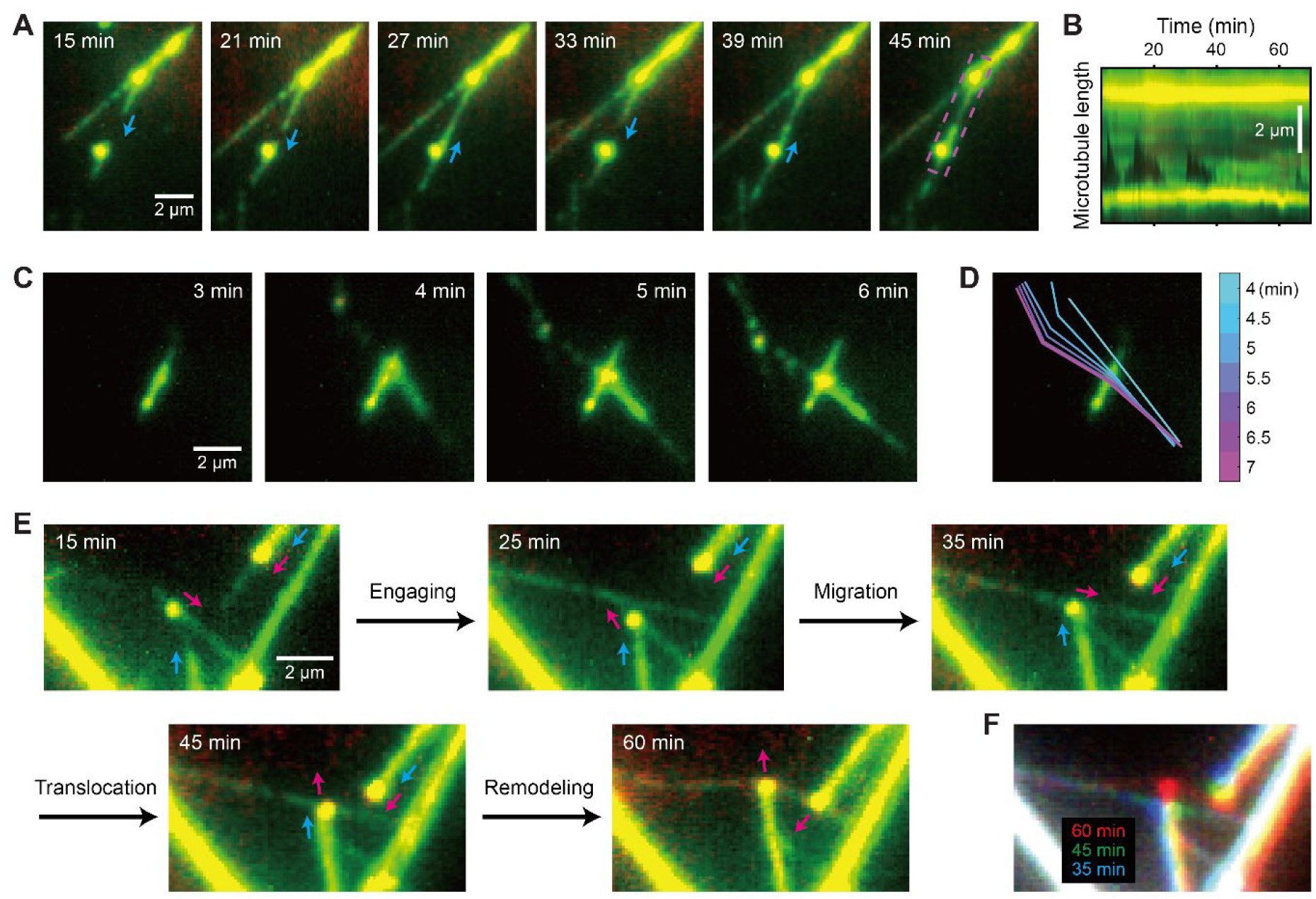
Microtubule network weaving and remodeling by tau-tubulin co-condensates. (A) Time-lapse images showing tau-tubulin co-condensates facilitating the longitudinal joining of microtubules, with blue arrows indicating the direction of microtubule extension. The pink dashed box marks the region analyzed for the kymograph in (B). (B) Kymograph illustrating the progressive association of microtubules over a 70-min period. (C–D) Lateral crosslinking of microtubules mediated by tau-tubulin co-condensates. (D) Trajectory plot showing points of intersection and crosslinking over time. (E) Sequential images capturing the migration of tau-tubulin co-condensates along microtubules. Blue arrows indicate microtubule growth, and red arrows indicate condensate movement, showing how interactions between growing microtubules and moving condensates contribute to structural remodeling. (F) Composite image showing microtubule deformation between 35 and 60 min, highlighting structural adjustments facilitated by tau-tubulin co-condensates.

Strikingly, tau-tubulin co-condensates further remodel the network by translocating along microtubule filaments and exerting mechanical forces. As they migrate using the filaments as pathways, they push against existing structures, causing bending and rewiring (Fig. 3E, Movie S8). Despite the inherent stiffness of microtubules, this movement results in significant adjustments (Fig. 3F), suggesting that co-condensates may target weak links or defects within the network. Collectively, these behaviors underscore their active role in dynamically reshaping the microtubule architecture.

Overall, the dynamic nature of tau-tubulin co-condensates contributes to microtubule network formation, not merely by nucleating and stabilizing microtubules but also by connecting existing structures and flexibly modifying configurations. Rather than being fixed, the network is dynamically woven over time, with the movement of tau-tubulin co-condensates leading to continuous remodeling, indicating that the network’s geometry evolves as they interact within the reticular structure.

### Voronoi tessellation for the analysis of tau clustering and tubulin enrichment

Since our observations of the early stages of microtubule organization revealed a central structuring role for tau-tubulin co-condensates, we next aimed to analyze how these condensates contribute to the final stabilized structures. To achieve this, we sought to interpret the in vitro network quantitatively by mapping tau and tubulin foci using Voronoi tessellation, a robust computational geometry technique that partitions an area into distinct regions based on proximity. In this method, each region or “cell” represents the area closest to its corresponding “seed” or reference point (in our case, the locations of tau foci), enabling intuitive yet precise spatial visualization. Given that this approach is well-suited for super-resolution microscopy because it effectively segments and quantifies protein clusters^20^, we reasoned that the Voronoi diagram would be practical for detecting tau clusters and mapping their influences on tubulin enrichment.

To localize tau and tubulin molecules, we used direct stochastic optical reconstruction microscopy (dSTORM), which extracts their positions from the blinking of fluorescent labels (Fig. 4A) (via the ThunderSTORM ImageJ plugin^25^). High-intensity spots of tau molecules, identified by blinking events above a set threshold, were then used as seed points for constructing Voronoi diagrams (using the function ‘voronoiDiagram’ in MATLAB). This approach allowed us to segment the full field of view of fluorescent images into numerous cells of varying sizes, each centered around tau locations for further analysis (Fig. 4D). In general, regions with more tau molecules are divided into more cells with smaller areas (see Fig. 4B, C, E, and F for more zoomed-in views), and thus the cell area itself serves as a first-hand proxy for determining the locations of tau clusters. Interestingly, we were able to distinguish at least three different log-normally distributed populations, each corresponding to the high-, medium-, and low-density regions (Fig. 4G).

**Figure 4.**
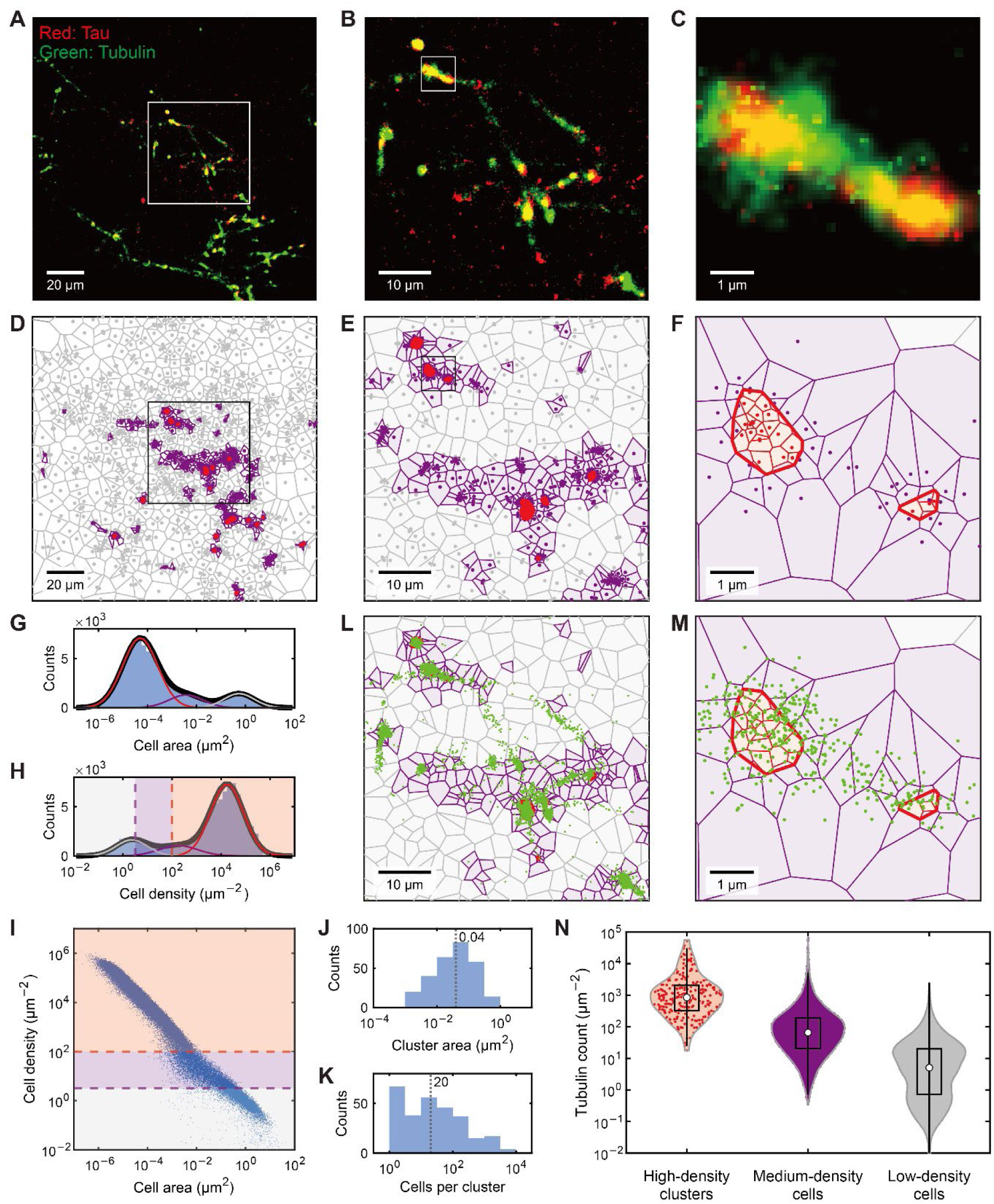
Voronoi tessellation for analyzing tau clustering and tubulin enrichment. (A–C) dSTORM images showing tau (red) and tubulin (green) distribution in microtubule networks. Panels (B) and (C) display zoomed-in regions from the white boxes in (A) and (B), respectively. (D–F) Voronoi tessellation used to segment tau foci, corresponding to views in (A–C). High-density tau cells are shown in red, medium-density cells in purple, and low-density cells in gray. In (F), high-density Voronoi cells consolidated into singular clusters are outlined with thick red borders. (G) Histogram of cell area distribution, revealing distinct populations based on tau density. (H–I) Histogram of cell density (H) and scatter plot showing the inverse correlation between cell area and density (I), confirming the density classification. Thresholds used for density-based classification are indicated by dashed lines. (J–K) Histograms showing cluster area distribution (J) and cell counts per cluster (K), indicating heterogeneity in cluster size. Median values are displayed. (L–M) Voronoi-based colocalization analysis of tubulin counts (green) within tau-defined regions. High-density tau clusters (outlined in red) show significant tubulin enrichment. (N) Violin plot of tubulin count within high-, medium-, and low-density tau-defined cells, revealing a positive correlation between tubulin enrichment and tau density.

To further verify tau clustering, we applied first-rank density analysis, a robust method previously used for detecting clusters in super-resolution images using Voronoi diagrams^20,21^. First-rank density, defined as the density of neighboring cells, was calculated for each Voronoi cell by dividing the number of directly adjacent cells by their total area. In larger, denser tau foci, tau events were more concentrated, leading to higher first-rank densities. The histogram for cell density (Fig. 4H) revealed distinct populations with proportions similar to those in the cell area distributions (Fig. 4G) and indeed showed a strong inverse correlation with cell area (Fig. 4I), confirming the validity of this approach. We accordingly classified the cells into three density groups across the tau dataset (Fig. 4D-F): high-density cells (> 10^2^ μm^−2^; red), medium-density cells (> 10^0.5^ μm^−2^; purple), and the remaining lower-density cells. To represent large, dense tau clusters as single objects, we consolidated contiguous high-density Voronoi cells into singular clusters, indicated by thick red borders in Fig. 4F. These clusters were heterogeneous in size (Fig. 4J, K), ranging from 10^−3^ to 1 μm^2^ per cluster, with a median of 0.04 μm^2^, implying a characteristic length scale of (0.04 μm^2^)^1/2^ = 0.2 μm.

Subsequently, to evaluate tubulin enrichment around tau foci, we measured the colocalization rate by counting tubulin blinking events as a function of tau density (Fig. 4L, M). Due to their extremely small areas, high-density Voronoi cells were not suitable for colocalization estimation, as they often resulted in empty tubulin counts. Therefore, we used the consolidated high-density clusters to estimate colocalization density around tau condensates. For comparison, we also measured tubulin counts within medium- and low-density Voronoi cells as defined by tau distribution. Our analysis revealed a clear correlation between tubulin enrichment and local tau density (Fig. 4N): tubulin counts were significantly higher within high-density tau clusters than in lower-density Voronoi cells. Medium-density cells, primarily situated at the edges of high-density clusters, showed intermediate tubulin counts, reflecting a gradual transition in tubulin concentration from tau condensates to the surrounding dilute phase. These findings suggest that tau clusters create localized regions of tubulin enrichment, supporting a role for tau in organizing microtubule assembly by concentrating tubulin at specific sites through a density-dependent recruitment mechanism. This density-based colocalization, confirmed by Voronoi analysis, underscores the active structural and organizational role tau clusters play in shaping the microtubule network.

### 3D Voronoi analysis of tau clustering and tubulin enrichment

Following our 2D analysis, we extended the Voronoi tessellation approach to three-dimensional space, incorporating depth to better capture spatial relationships between tau clusters and tubulin within microtubule networks. Due to technical limitations, confocal microscopy was employed to obtain z-stack images, and these images were processed differently to produce tau and tubulin coordinates (see Methods). Although confocal imaging offers lower resolution than dSTORM, this approach effectively identified tau cluster locations across the full z-stack. Using 3D reconstructions of confocal fluorescence images (Fig. 5A–C, Movie S9), we segmented tau images into Voronoi cell volumes, categorizing regions of tau density as high, medium, or low (colored red, purple, and gray in Fig. 5D–F). This segmentation enabled analysis of tau distribution and its impact on tubulin localization in 3D, providing a comprehensive view of tau-tubulin interactions within microtubule networks.

**Figure 5.**
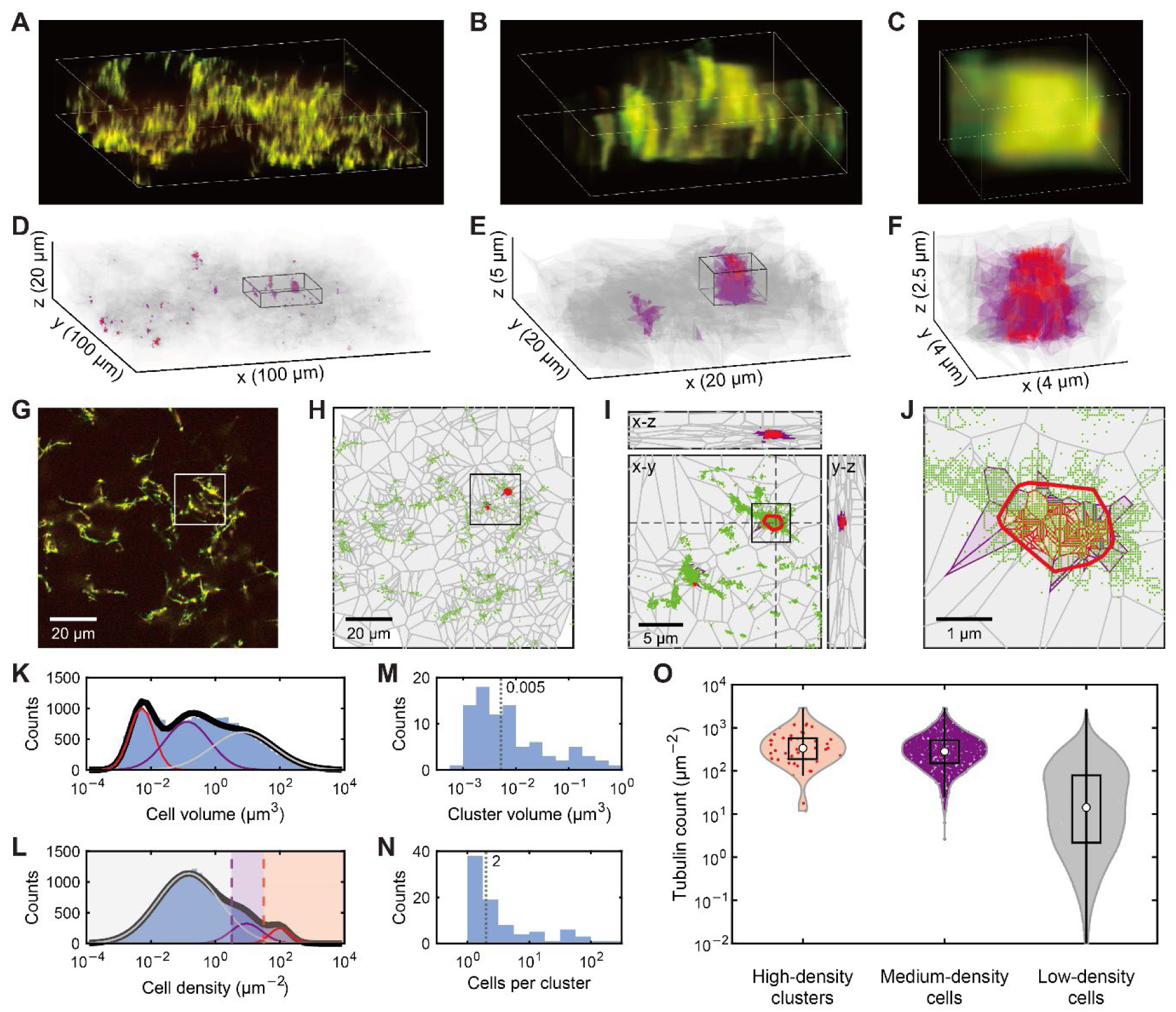
3D Voronoi diagram analysis of tau-tubulin clustering and colocalization. (A–C) 3D reconstructions of confocal fluorescence images showing tau (red) and tubulin (green) within microtubule networks. (D–F) 3D Voronoi cell segmentation of tau clusters from images in (A–C). High-density tau cells are shown in red, medium-density in purple, and low-density in gray. Boxed regions in (D) and (E) indicate areas for zoomed-in views in (E) and (F), respectively. (G–J) Representative x-y plane image from the 3D confocal stack (G) and cross-sections of 3D Voronoi cells from tau foci (H–J), with overlaid tubulin localization (green). Boxes show zoomed-in regions. In (I), cross-sectional views of a high-density tau cluster (outlined in thick red) are shown in x-y, x-z, and y-z planes to illustrate the spatial extent and morphology of tau-tubulin clusters. (K–L) Histograms of cell volume (K) and cell density (L) distributions, showing distinct populations based on tau density in 3D. Dashed lines mark thresholds for density-based classification. (M–N) Histograms of the volume distribution of 3D tau clusters (M) and the number of Voronoi cells per cluster (N), indicating cluster heterogeneity. Median values are shown. (O) Violin plot of tubulin counts within high-, medium-, and low-density tau-defined regions, showing a positive correlation between tubulin enrichment and tau density in 3D.

To better visualize the spatial organization of 3D tau clusters and tubulin enrichment, we examined x-y, x-z, and y-z cross-sectional views of the Voronoi diagrams corresponding to a representative confocal image slice (Fig. 5G–J, Movie S10). These views revealed that medium- to high-density tau cells form local clusters, consistent with the 2D analysis. The distributions of Voronoi cell volumes and densities further supported the classification of tau regions into high-, medium-, and low-density populations (Fig. 5K, L). Distinct peaks in cell volume and density histograms corresponded to these density classes, indicating heterogeneity in the 3D organization of tau clusters. Similar to the 2D approach, we clustered high-density cells by consolidating neighboring volumes. Notably, these clusters (outlined in red in Fig. 5I, J) frequently colocalized with regions of elevated tubulin fluorescence, again consistent with the 2D findings. The distribution of cluster volumes (Fig. 5M) and the number of Voronoi cells per cluster (Fig. 5N) revealed substantial variability in cluster size and composition, with larger clusters generally comprising multiple high-density cells, suggesting a hierarchical organization of tau clustering. Despite methodological differences between 2D and 3D analyses, the resulting median volume of high-density tau clusters (0.005 μm^3^), with a length scale of (0.005 μm^3^)^1/3^ ≈ 0.17 μm, aligns closely with the 2D cluster size of 0.2 μm.

Finally, to assess tubulin enrichment within tau-defined regions of varying density, we measured tubulin counts in each region (high, medium, and low density) and visualized the results in a violin plot (Fig. 5O). Our analysis revealed a positive correlation between tau density and tubulin enrichment, with high-density tau clusters exhibiting the highest tubulin counts. The distinction between medium- and high-density regions was less clear, likely due to the limited resolution of confocal microscopy, which barely resolved the high-density cells. Nonetheless, low-density tau regions consistently showed the lowest tubulin enrichment. This trend aligns with our 2D findings, reinforcing the role of tau clusters in creating localized areas of tubulin enrichment and supporting the assembly and organization of the microtubule network through density-dependent recruitment.

## Discussion

Our study reveals novel roles for tau-tubulin interplay in microtubule assembly and organization, extending the known function of tau in promoting local tubulin enrichment for microtubule nucleation^11^. Traditionally, tau has been viewed primarily as a stabilizer of microtubule structures, often acting on microtubules in droplet form^26,27^, with limited focus on its role in recruiting tubulin as a client molecule within tau condensates. Our findings shift this perspective, highlighting tau-tubulin co-condensates as active structuring elements of the microtubule network. Real-time imaging (Figs. 2, 3) demonstrated that these co-condensates influence microtubule dynamics through their liquid-like properties, enabling both nucleation and continuous remodeling.

Furthermore, using Voronoi diagram analysis (Figs. 4, 5), we defined areas of influence around tau and assessed tau-tubulin colocalization, revealing that larger and denser tau clusters enhance local tubulin accumulation, thereby promoting co-condensate formation and subsequent microtubule nucleation. This dual activity—stabilizing nascent filaments and reshaping the network—highlights their critical role in dynamically organizing and maintaining the microtubule cytoskeleton.

Building on these insights, we propose a dynamic weaving model (Fig. 6) in which tau-tubulin co-condensates continuously interact with and remodel microtubules through mechanisms such as bridging, bundling, crosslinking, diffusion, and merging. By connecting microtubule ends and forming crosslinks, they create a flexible, reticular architecture that adapts over time. Their mobility allows them to move along microtubules, merge with other condensates, and induce structural adjustments, contributing to a dynamic, woven network. This model positions tau-tubulin co-condensates as central to both the stabilization and adaptability of the microtubule cytoskeleton, allowing it to respond to cellular needs and environmental cues.

**Figure 6.**
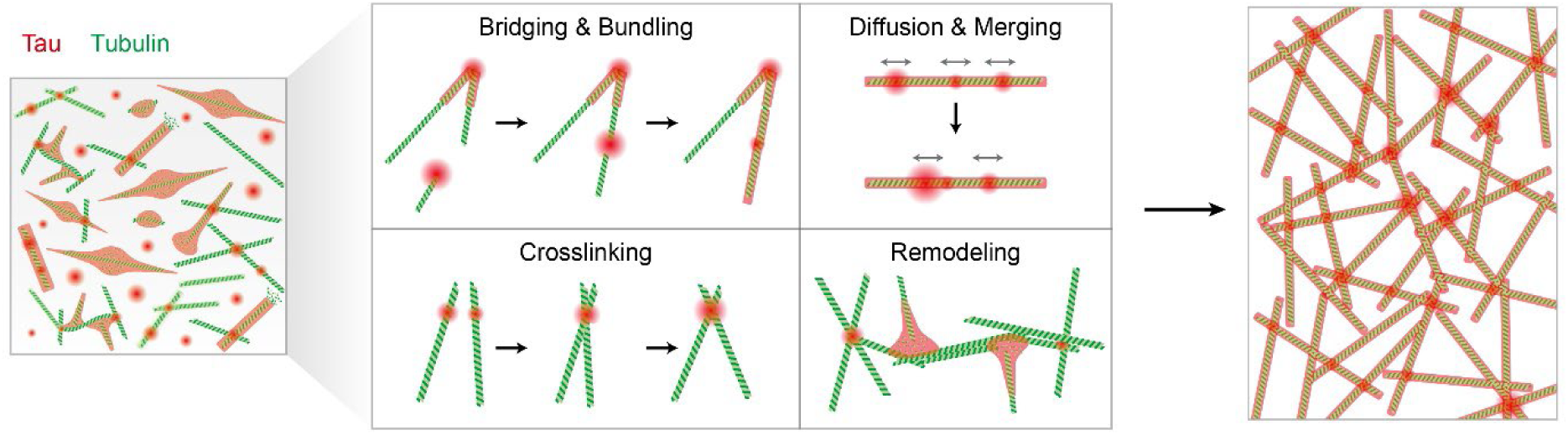
Dynamic weaving model for the microtubule structuring by tau-tubulin co-condensates. Illustration of tau-tubulin co-condensates dynamically organizing microtubules through distinct mechanisms, including: (i) bridging and bundling, where co-condensates connect and align microtubule ends; (ii) crosslinking, where co-condensates link intersecting microtubules to enhance network stability; (iii) diffusion and merging, where co-condensates move along microtubules and merge to form larger clusters; and (iv) remodeling, where co-condensates induce structural changes by exerting force on microtubules. The resulting organized network, formed through continuous tau-tubulin interactions, illustrates a woven, reticular architecture. This network can be further stabilized by tau spreading onto the surfaces of microtubules as the structure reaches equilibrium.

By systematically examining the tau/tubulin/PEG system across varying component concentrations, we identified fine-tuned, physiologically relevant conditions that support successful microtubule polymerization without requiring unnaturally high protein levels or excessive crowding agents (Fig. 1). Our phase diagrams revealed an optimal range of tau, tubulin, and PEG concentrations that yield well-formed microtubules and their reticular network. These findings provide valuable guidelines for future experiments, enabling precise adjustments of parameters to achieve specific outcomes, as shown with the Voronoi analysis. The results further suggest that striking a balance in tau and tubulin concentrations may be crucial in cells to maintain both structural stability and dynamic flexibility. It is tempting to speculate that this balance could be further fine-tuned by cellular mechanisms, such as tau phosphorylation, to regulate co-condensate dynamics in response to cellular needs.

The co-condensates can deform existing microtubule structures by converting the force from microtubule growth into mechanical work (Fig. 3E, F). This deformation is particularly notable given the intrinsic stiffness of microtubules, which generally resist bending on short length scales. Since tau has been shown to preferentially associate with curved regions of microtubules to stabilize them^27^, our observations once again highlight a seemingly contradictory effect of tau condensates on microtubule stability. The local deformation of microtubules driven by tau-tubulin co-condensates can potentially involve two mechanisms. First, these co-condensates could locally “melt” microtubules by dissolving tubulin on their surface, with a slight mechanical “push” assisting the process^28^. Alternatively, co-condensates may preferentially target weak points in the microtubule network, such as areas with defects, structural discontinuities, or regions where the number of filaments within bundled microtubules is inconsistent. These weak links could serve as focal points for tau-tubulin co-condensates to exert mechanical forces, leading to localized bending and remodeling. In either case, the presence of tau-tubulin co-condensates at these points of force application acts like molecular solder. They protect the microtubule network from complete rupture by stably holding the disjoined filaments together while allowing slight modifications. It would be of interest to use single-molecule force techniques to study the interplay between the applied mechanical load and co-condensate activity.

Extending this study to cell imaging, particularly during the early phases of microtubule assembly, could reveal how these co-condensates function in the natural cellular environment. For instance, after chemically disrupting microtubules using agents like nocodazole, we could monitor the rebuilding of the microtubule network to assess the involvement of tau-tubulin co-condensates in nucleation and organization. Additionally, examining microtubules during and after cell division, when the cytoskeleton undergoes significant remodeling, could further elucidate the mechanisms by which tau-tubulin co-condensates contribute to microtubule dynamics in vivo. Such live-cell studies would bridge the gap between our in vitro observations and the complex behavior of microtubules in living systems.

The Voronoi approach used here can be extended to other cytoskeletal network analyses that require examination on multiple length scales. For instance, similar methods could be applied to study actin filament formation by analyzing the colocalization of actin and its nucleating factors, such as the Arp2/3 complex or formins^29^. Another example includes mapping microtubule nucleation mediated by the targeting protein for Xklp2 (TPX2) when it forms co-condensates with tubulins^30^. Additionally, this approach could be applied to the formation of mitotic spindles facilitated by γ-tubulin ring complexes (γ-TuRCs) during cell division^31^. Such applications would help us understand how localized patterns, such as protein clustering or signaling events, influence the overall structure and dynamics of the cytoskeletal network through a mathematically concise and simplified mapping. Interestingly, we have recently discovered in vitro that tau forms dynamic condensates on naked DNA strands, which further recruit microtubules to form complex networks^32^. It would be intriguing to explore whether the Voronoi-based analysis could be applied to this DNA-tau-microtubule system to reveal spatial patterns and organization principles.

## Methods

### Tau purification and labeling

Full-length human tau protein with a 6x His-tag was cloned into the pRK172 vector and expressed in LEMO21 cells (NEB #C2528J). Cells were cultured in TB media at 37 °C until the OD600 reached 0.6, at which point 10 mM betaine and 500 mM NaCl were added. The culture was then maintained at 30°C for 30 min. Subsequently, 0.4 mM IPTG was introduced, and culturing continued for an additional 3–4 h. Cells were harvested by centrifugation at 4 °C and 3000 g for 20 min, resulting in pellets. These pellets were resuspended in lysis buffer (20 mM MES pH 6.0, 0.2 mM MgCl_2_, 1 mM EGTA, 1X PMSF, 1X protease inhibitor cocktail, 5 mM DTT) and homogenized. The cells were sonicated at 30% amplitude with 30 s on-off cycles for 3 min, immediately followed by the addition of 500 mM NaCl. The lysate was then boiled for 15 min and centrifuged at 15,000 g for 1 h at 4 °C. The resulting supernatant was purified using Ni-NTA resin. The bound resins were washed with wash buffer (50 mM phosphate buffer pH 7.2, 200 mM NaCl, 20 mM imidazole) and then eluted with elution buffer (50 mM phosphate buffer pH 7.2, 200 mM NaCl, 500 mM imidazole). Custom-made TEV protease was added to remove the histidine tag, followed by overnight incubation at 4 °C. Additional purification was achieved through ion exchange chromatography (Bio-rad #1580030) to remove dimers and other undesired proteins. The purified proteins were then concentrated using a protein concentrator (Sartorius #VS15T01) and verified by SDS-PAGE (Fig. S1). Protein labeling was achieved by mixing the protein with Alexa Fluor 488-maleimide (Sigma, #A10254) at a 20:1 molar ratio of fluorescence-tagged to wild type protein. The mixture was incubated for 2 h at room temperature with gentle mixing in a reaction buffer (50 mM phosphate buffer, 0.05 mM TCEP). After removing the free dye molecules by gel filtration (Bio-rad #1500738), the dye-labeled protein was concentrated and stored until use.

### Microtubule formation by tau-tubulin co-condensation using molecular crowding agents

Tau droplets were pre-formed under two distinct conditions, each incorporating 5–10% Alexa 488-labeled tau: 5 μM tau with 7.5% PEG or 25 μM tau with 10% dextran, unless otherwise noted. Both conditions used BRB80 buffer (80 mM PIPES, 1 mM EGTA, 1 mM MgCl2, pH 6.9) supplemented with 100 mM NaCl, 100 nM PCD, and 5 mM PCA. After 1 h incubation at room temperature, 5 μM of porcine tubulin dimers (Cytoskeleton #T240-B), including 5% HiLyte 647-labeled tubulin dimers (Cytoskeleton #TL670M-A), were introduced into the tau droplets along with 1 mM GTP and 1 mM DTT. These mixtures were then injected into a PEGylated glass chamber, composed of quartz glass and 60 mm long coverslips as previously described^33^. After injection, the incubation time of the samples varied depending on the experimental objectives. For dSTORM and confocal imaging, samples were imaged after incubating 2 h at room temperature. However, for nucleation experiments, imaging was conducted throughout the incubation period to monitor microtubule formation in real time. To prevent evaporation and maintain sample integrity, both the inlet and outlet of the chamber were sealed with epoxy.

### Optical setup

A custom-built optical setup was integrated into an inverted microscope (Olympus IX73), including a 60× oil-immersion lens and an OptoSplit II ByPass (CAIRN Research) for dual-color fluorescence imaging. The setup employed continuous lasers (Cobolt) with wavelengths of 488 nm and 633 nm to excite Alexa 488-labeled tau and HiLyte 647-labeled tubulin, respectively.

To enable simultaneous visualization of both fluorophores, the two excitation beams were co-aligned using a 525 nm dichroic mirror, magnified tenfold with a 10X expander, and focused onto the sample through an achromatic lens. The emitted fluorescence was captured through a 405/488/532/635 nm quad-edge dichroic mirror (Semrock) and then separated by the OptoSplit II ByPass using a 614 nm dichroic mirror. The fluorescence was filtered through a 708(75) nm bandpass filter for tubulin and a 512(25) nm bandpass filter for tau. Images were acquired by an sCMOS camera (Teledyne Photometrics, Prime BSI Express) typically with 50 ms resolution and a ∼50-μm field of view.

### dSTORM and confocal imaging of tau/tubulin/PEG system

To conduct 2D dSTORM imaging, we introduced 142 mM β-mercaptoethanol and 2 mM cyclooctatetraene into the tau/tubulin/PEG mixture during the microtubule formation. For illumination, the tubulin channel was illuminated at an intensity of 36 W/mm^2^, while the tau channel was illuminated at 12 mW/mm^2^. For each dSTORM image reconstruction, 10,000 frames were recorded with an exposure time of 50 ms per frame at a pixel size of 108 nm. Single-molecule localization was performed using the thunderSTORM plugin in ImageJ. For 3D confocal imaging, the sample preparation followed the protocol described in the microtubule formation section. Imaging was performed using a Zeiss LSM 900 confocal microscope equipped with Airyscan 2, conducting z-stacking with a 240 nm step size to capture the three-dimensional structure of the microtubule network at a pixel size of 43 nm. A total of 100-200 z-section images were acquired and subsequently reconstructed using ZEN blue 3.4 software.

### Voronoi diagram and colocalization analysis

To construct the Voronoi diagrams for the tau structures, the dSTORM results and confocal fluorescence images were processed utilizing custom MATLAB codes (https://github.com/ShonLab). In 2D dSTORM-based analysis, the x,y-coordinates of single-molecule localization were directly fed to the MATLAB functions delaunayTriangulation and voronoiDiagram to construct Voronoi diagrams, after excluding low-intensity peaks. In 3D confocal-based analysis, the images were first thresholded to identify high-intensity regions, and then the MATLAB function bwmorph was applied to iteratively shrink connected components in the binary images, yielding ultimately eroded points at the locations of tau clusters. For both 2D and 3D, the resulting Voronoi cells were classified into three groups based on their first rank density of neighboring cells. Additionally, the neighboring high-density cells were consolidated into clusters for analyzing colocalization. Local features, including the area/volume of each region and the count of tubulin fluorophores, were then calculated.

## Author Information

## Acknowledgments

We thank Sang Hyeok Jeong, Jaehyeon Shin and Haeun Yoo for the help with experiments and analysis; and the members of the Shon laboratory for discussions and assistance. This work was supported by the National Research Foundation of Korea (NRF) grant funded by the Korea government (MSIT) (NRF-2022R1C1C1012176, RS-2023-00218927, and RS-2024-00344154).

## Author Contributions

C.L.-E. conceived the project. C.L.-E., J.J., and M.J.S. designed the project. C.L.-E., J.J., and C.P. prepared samples. C.L.-E. and J.J. performed fluorescence imaging. C.L.-E. and M.J.S. analyzed data. All authors prepared the figures. C.L.-E. and J.J. wrote the first draft and M.J.S. edited the manuscript.

