## Supplementary Information for "Structuring Role of Tau-Tubulin Co-Condensates in Early Microtubule Organization"

Lee-Eom *et al.*

### Supplementary Figures

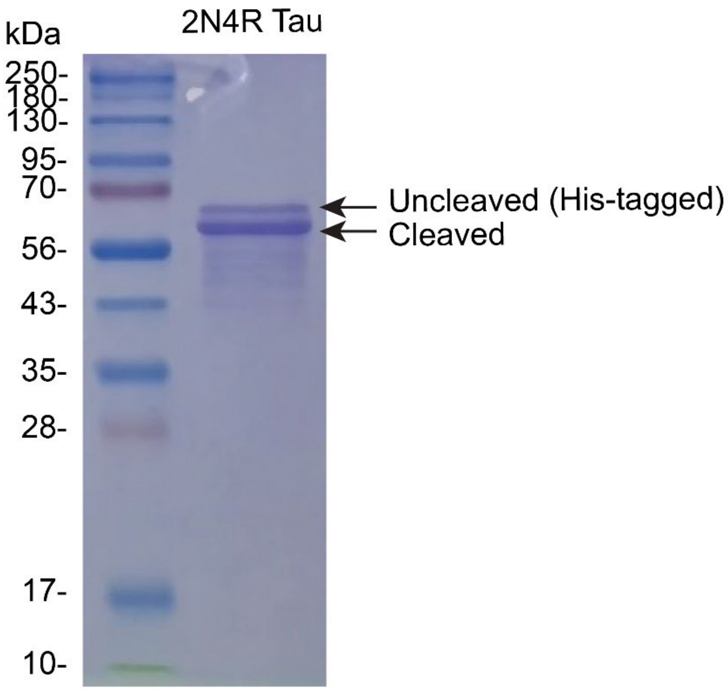


**Figure S1. Purified 2N4R tau protein**

SDS-PAGE gel showing purified 2N4R tau proteins. Bands indicate uncleaved (His-tagged) and cleaved forms of 2N4R tau protein.

### Supplementary Video Legends

For all movies, the red channel shows Alexa 488-labeled tau, and the green channel shows HiLyte 647-labeled tubulin. Exposure time was 0.2 s.

**Movie S1. Time lapse of microtubule nucleation**
A time-lapse video of real-time nucleation of microtubules with tau-tubulin co-condensates in the presence of 10% dextran. Snapshots from this video are presented in Fig. 2A of the main text. Frames are averaged over 10 s and the playback is sped up 300×.

**Movie S2. Rapid nucleation of microtubules**
A video of rapid nucleation of microtubules. Snapshots from this video are presented in Fig. 2B of the main text. Frames are averaged over 1 s and the playback is sped up 30×.

**Movie S3. Dynamic instability of microtubules with varying levels of tau**
A video of microtubule instability during nucleation. Snapshots from this video are presented in Fig. 2D of the main text. Frames are averaged over 1 s and the playback is sped up 30×.

**Movie S4. Diffusion of tau-tubulin co-condensates along microtubules**
A video of moving tau-tubulin co-condensates along microtubules. Snapshots from this video are presented in Fig. 2F of the main text. Frames are averaged over 10 s and the playback is sped up 300×.

**Movie S5. Merging of tau-tubulin co-condensates on microtubules**
A video of merging tau-tubulin co-condensates. Snapshots from this video are presented in Fig. 2H of the main text. Frames are averaged over 10 s and the playback is sped up 300×.

**Movie S6. Longitudinal joining of microtubules mediated by tau-tubulin co-condensates**
A video of microtubules longitudinally bridged and joined by tau-tubulin co-condensates. Snapshots from this video are presented in Fig. 3A of the main text. Frames are averaged over 10 s and the playback is sped up 300×.

**Movie S7. Lateral assembly of microtubules mediated by tau-tubulin co-condensates**
A video of microtubules laterally crosslinked by tau-tubulin co-condensates. Snapshots from this video are presented in Fig. 3C of the main text. Frames are averaged over 1 s and the playback is sped up 30×.

**Movie S8. Dynamic weaving of microtubules driven by tau-tubulin co-condensates**
A video of dynamic weaving of microtubules by tau-tubulin co-condensates. Snapshots from this video are presented in Fig. 3E of the main text. Frames are averaged over 10 s and the playback is sped up 300×.

**Movie S9. 3D Reconstruction of tau-tubulin network in PEG environment**
A 3D reconstruction of the tau-microtubule network, created from confocal microscopy z-stack images, as shown in Fig. 5 of the main text. The solution contained 5 μM tubulin and 5 μM tau in a 7.5% PEG environment.

**Movie S10. Z-stack visualization of 3D Voronoi diagrams for the tau-tubulin network**
Visualization of the Voronoi diagrams in x-y planes throughout the z-stack of confocal images, as shown in Fig. 5G, H of the main text.
